# Climate Gradients and Habitat Discontinuity Structure Genetic Variation in a Spring-Specialist Plant

**DOI:** 10.64898/2026.05.08.723645

**Authors:** Matthew Weiss, Trevor M. Faske, Liza M. Holeski

## Abstract

**Background and Aims:** Groundwater-dependent ecosystems support disproportionate biodiversity in arid regions, yet the population genetics of spring-specialist plants remains poorly understood. Here, we present the first species-wide genetic dataset for crimson monkeyflower ( *Mimulus verbenaceus*, Phrymaceae), a spring-specialist plant distributed in seeps, springs, and associated riparian areas across desert regions of North America.

**Methods:** Using genome-wide reduced representation sequencing data consisting of 10,760 SNPs from 175 individuals across 17 populations, we characterized the patterns of genetic diversity using *F*_ST_ and Nei’s *D.* Population structure was assessed using ADMIXTURE and PCA. We examined the contributions of climate to range-wide genetic variation in crimson monkeyflower using redundancy analysis.

**Key Results:** Patterns of genetic differentiation were more consistent with those of spring-specialist animal taxa than those of upland plants or generalist riparian plants. We found strong population structure at both broad regional scales and at fine local scales. While riparian connectivity influenced local patterns of diversity, adaptation to local climatic variation was more influential at regional scales, with temperature, relative humidity, and a monsoon-driven climate gradient structuring genetic differentiation.

**Conclusions:** Our findings highlight the distinctive influence of isolated perennial groundwater sources, as well as adaptation to climate, in shaping genetic variation in this spring-specialist plant. These findings suggest that spring-specialist plants deserve special consideration in ecological theory, management, and conservation.

## INTRODUCTION

Groundwater-dependent ecosystems such as seeps and springs play a disproportionate role in supporting biodiversity, acting as keystone ecosystems in arid regions (Cartwright and Johnson 2018; Nielson *et al*. 2019; Work 2023). These environments are frequently hydrologically decoupled from regional and short-term climate variation, serving as stable refugia from drought events that are becoming less predictable and more common due to climate change (McLaughlin *et al*. 2017). Yet, desert springs are increasingly under threat from groundwater depletion, habitat fragmentation, livestock, invasive species, and climate change itself (Kløve *et al*. 2014; Cartwright and Johnson 2018). As such, the rich organismal diversity adapted to these ecosystems is also under threat.

Organisms adapted to the unique environmental constraints of seeps and springs are referred to as crenophiles or spring-specialist species (Danks and Williams 1991; Smieja 2014; Jyväsjärvi *et al*. 2015). Much of the work in these species has focused on animal taxa, especially aquatic invertebrates (Jyväsjärvi *et al*. 2015; Stevens *et al*. 2020; Blattner *et al*. 2022; Violeta and Mateusz 2023). While spring-specialist plants provide crucial habitat for other species and are at risk from threats to their ecosystems, little is known about their genetic diversity, population structure, and patterns of local adaptation. Investigating the genetic variation and structure of spring-specialist plants can provide insights to these foundational species that other organisms in these ecosystems depend upon.

Explorations of population structure and genetic diversity in spring-specialist animal taxa have revealed patterns shaped by discrete patches of suitable habitat and ancient riparian connectivity (Houston *et al*. 2015; Goudarzi *et al*. 2019; Campbell *et al*. 2022). Spring-specialist plants may show patterns more similar to spring-specialist animal taxa than to upland plants or generalist riparian plants. Plant populations at smaller and more isolated springs are expected to experience different demographic processes than upland plants, such as reduced gene flow, increased genetic drift, and reduced within-population genetic diversity (Orsini *et al*. 2013). This pattern is expected to be particularly pronounced when springs serve as rare oases within largely arid regions (Houston *et al*. 2015; Campbell *et al*. 2022). Alternatively, since springs are frequently found in riparian corridors, spring-specialist plants might show patterns of diversity seen in other riparian species if their seeds or propagules are adapted for hydrochory. If riparian connectivity plays an important role for spring-specialist plants, we may expect to see greater diversity across populations connected by riparian corridors (Bothwell *et al*. 2017) and higher genetic diversity in upstream populations than downstream populations (Ritland 1989; Honnay *et al*. 2010).

The genetics of spring-adapted plants may also be influenced by selection from different climate gradients when compared to upland plants, especially in arid regions. In particular, local adaptation to precipitation and drought is often an important driver of genetic differentiation across species of plants in arid regions (Li *et al*. 2018; Faske *et al*. 2021; Omondi *et al*. 2024; Shryock *et al*. 2024). For spring-specialist species, the availability of groundwater is decoupled from short-term and extreme precipitation events and may buffer selective pressure on drought adaptation. As a result, climate variables other than precipitation may play a more prominent role in shaping genetic structure in these species.

Crimson monkeyflower (*Mimulus verbenaceus*, Phrymaceae) presents an ideal system for studying connectivity and diversity within a spring-specialist plant. Across its range, crimson monkeyflower occupies seeps, springs, and associated perennial riparian areas from canyon bottoms to isolated mountain ranges in deserts of Western North America (Beardsley *et al*. 2003; Vickery 2008). The perennial water sources in this arid region are particularly threatened and vulnerable to environmental change (Castle *et al*. 2014; Stevens *et al*. 2020). Within these habitats, crimson monkeyflower plays an important role as a food source for local and migratory hummingbirds and provides habitat for other animals such as the critically endangered Kanab ambersnail (Vickery 1992; Vickery and Sutherland 1994; Spamer and Bogan 1998) . Our study will provide a baseline understanding of one of the foundational species in these systems, which could act as a crucial tool for future monitoring efforts.

Crimson monkeyflower is a model system for research in novel trait evolution, trait syndromes, and functional genetics (Yuan 2019; Chen *et al*. 2022; LaFountain *et al*. 2022; Wenzell *et al*. 2025). Despite growing interest in this species, little is known about its natural ecology, evolution, and population genetics, including levels of genetic diversity and population structure across its range. By characterizing how demographic processes and environmental gradients shape population structure in crimson monkeyflower, this study provides a foundation for connecting functional genetic research to species-wide processes shaping trait evolution in this model system.

To assess the factors influencing genetic differentiation across the range of crimson monkeyflower, we generated and analyzed the first range-wide genetic dataset for this species. Specifically, our goals are to (1) assess levels of genetic diversity within and among populations and characterize patterns of population structure both across the species range and at fine local scales, (2) understand the contributions of both climate and geography to species-wide genetic variation, and (3) quantify the proportion of genetic variation associated with specific climate variables driving local adaptation.

## MATERIALS AND METHODS

### Sample Collection and Library Preparation

Crimson monkeyflower grows across a largely arid region of western North America, including the Sonoran Desert. Populations are largely limited to relatively wetter and cooler landscape features within this region (Beardsley *et al*. 2003; Vickery 2008; Sheth and Angert 2014) . These landscape features include canyon bottoms associated with riparian corridors and high-elevation mountain ranges. High-elevation features within this region include the Sierra Madre Occidental in the southern part of the range and the Colorado Plateau to the north. A chain of mountains, the Madrean Sky Islands, connects these two large high-elevation regions in the center of our species’ range, creating habitable pockets for our species surrounded by otherwise uninhabited lowlands (López-Hoffman and Quijada-Mascareñas 2012) . Evidence from sediment cores and packrat middens shows that current distributions of vegetation in these sky islands represent refugia during Holocene drying and are remnants of previously more connected lowland ecosystems (Baker 2008; Galaz-Samaniego *et al*. 2023).

We collected tissue from plants from 17 populations of crimson monkeyflower from the Southwestern United States. Crimson monkeyflower populations are limited to landscape features that support perennial water sources, such as mountain ranges and riparian corridors. Our populations were collected from eight major landscape features, including two mountain ranges (Huachuca Mountains and Black Mountains) and six riparian corridors (Gila River, Aravaipa Canyon, Lower Verde River, Oak Creek, Grand Canyon and Virgin River). For each population, between four and ten individuals were sampled for genetic analysis. We obtained 19 additional tissue samples from herbaria specimens, including six samples from a morphologically distinct lineage near Durango, Mexico, resulting in a total of 189 samples. Tissue collections consisted of 2-3 leaves, which we dried in paper envelopes using silica gel. We ground tissue using a FastPrep-24 homogenizer (MP Biomedicals) and extracted DNA from samples using a modified CTAB protocol (Doyle and Doyle 1987)with quality control performed using nanodrop and gel electrophoresis. We digested genomic DNA samples using EcoRI and MspI before ligating to adapters and indices as part of a reduced representation protocol (ddRADseq; Peterson et al., 2012) . We performed size selection using a Pippin Prep (Sage Science, Beverly, Massachusetts, USA) to select fragments between 400 and 600 bp. Samples were then pooled into a single library and sequenced at the Genomics and Cell Characterization Core Facility at the University of Oregon using a single lane of an Illumina NovaSeq 6000, SP 100 cycle run.

### Alignment and Genotyping

We demultiplexed sequences using fastq-multx (Aronesty 2013) and completed sequence assembly and genotype calling using ipyrad (Eaton and Overcast 2020). Raw reads were quality filtered to allow a maximum of 5 low-quality bases per read (Q<20) and require a minimum read length of 35 bp after adapter trimming. Adapter sequences were removed, and reads were mapped to the *Mimulus verbenaceus* genome (MvBLg_v2.0) (LaFountain *et al*. 2022) using a clustering threshold of 85% similarity. Consensus base calls required a minimum depth of 6 reads (both for statistical and majority-rule calling), with clusters exceeding 10,000 reads excluded. The datatype for assembly was specified as ‘ddrad’ for this assembly, with two restriction overhangs specified (AATTC, CGG). Initial variant calling produced a dataset of 150,710 loci across 189 individuals. We used VCFtools to filter this dataset to remove non-biallelic loci. Loci were further filtered to retain only those with a minor allele frequency (MAF) ≤ 0.02 and to remove those with ≥ 60% missing data. Individuals were then filtered to exclude those with > 60% of their loci missing. Lastly, loci were thinned to one SNP per tag to minimize linkage among markers.

### Genetic Diversity and Population Structure

To estimate genetic structure within crimson monkeyflower, we used both an ordination-based method (principal component analysis [PCA]) and a model-based maximum likelihood method (ADMIXTURE). Except where noted, all downstream analyses were performed in R v4.3.3 (R Core Team 2020). A PCA was run across all sampled individuals and eigenvalues were extracted to calculate percent variance explained for each principal component (PC). Ancestry proportions for each individual were estimated using a model-based clustering algorithm implemented in ADMIXTURE v1.3.0 (Alexander *et al*. 2009). We then ran ADMIXTURE for values of K (number of population clusters) ranging from 1 to 10 with 10-fold cross-validation to assess model fit. The optimal K was then determined using the lowest cross-validation error and through assessing ΔK in a manner similar to the method described in Evanno et al., 2005. To visualize patterns of genetic clustering across the landscape, we plotted individuals based on geography and their population clusters as determined by ADMIXTURE analysis.

To examine fine scale population structure, we conducted a PCA on two subsets of our samples from well sampled landscape features. One PCA included 69 individuals from seven sampling locations within a well connected riparian network, whereas the other included 29 individuals from sampling locations lacking connectivity via perennial water within an isolated mountain range.

For population genetic analysis, populations represented by four or fewer individuals were excluded to ensure reliable estimates of allele frequencies and to reduce bias in downstream analyses. We retained 16 populations of crimson monkeyflower for population-level analyses, each represented by 7–10 individuals after filtering (total n = 155). Unequal sample sizes (7–10 individuals per population) were accommodated by applying sample-size weighting or bootstrapped resampling where appropriate. The resulting data set represented sites across the species’ range within the southwestern United States. Sampling locations correspond to distinct isolated populations frequently associated with a single seep or spring. To estimate genetic diversity within populations, we calculated expected heterozygosity (He). To quantify pairwise differentiation between populations, we calculated *F*_ST_ (Weir and Cockerham 1984) and Nei’s genetic distance (Nei 1972).

### Isolation by Distance and Adaptation to Climate Gradients

Climate variables for each site were obtained from ClimateNA (Wang *et al*. 2016) using latitude, longitude, and elevations obtained from the Elevation Point Query Service (U.S. Geological Survey 2019) using the elevatr package (Hollister *et al*. 2023). We then obtained 30-year climate normals from 1991-2020 for a total of 25 climate variables for subsequent assessments of the influence of climate variation on genetic differentiation. We reduced the dimensionality of the dataset using a principal component analysis (PCA) on scaled climate variables (mean =0, SD = 1). The first three dimensions of climate variation (Dim1, Dim2, Dim3) were retained for downstream analysis.

To assess demographic and adaptive processes driving genetic structure, we conducted tests of IBD and IBE by assessing the effects of geographic and climate distance on genetic differentiation among our populations. To assess IBD, we calculated pairwise Haversine distance between sites using geosphere, v. 1.6-5. To assess IBE, we calculated Euclidean distance between sites using PCA scores from the first five dimensions of climate variation described above. The relationship between both geographic and climate distance and genetic distance (linearized *F*_ST_; *F*_ST_ / (1-*F*_ST_) ) was estimated using Mantel tests (Mantel 1967) implemented in vegan. Additionally, we assessed the relationship between genetic distance and climate distance when controlling for geographic distance using a partial Mantel test. Pearson correlation coefficients were used to assess the strength of the association between distances, while statistical significance (alpha > 0.05) was assessed using 9999 random permutations.

To evaluate how specific climate gradients contribute to the spatial genetic structure across the landscape, we used a redundancy analysis (RDA) in the vegan package. The first three dimensions of climate variation (described above) were used as predictors in our RDA with genome-wide genetic variation as the response variable. We computed constrained axes to quantify the variation in the genetic dataset predicted by climate variation. Adjusted R² was calculated by comparing constrained and unconstrained models, representing the total proportion of genetic variation explained by adaptation to climate. Significance of the overall model and individual RDA axes was assessed using permutation tests. To evaluate the unique contribution of each dimension of climate variation, we conducted partial redundancy analysis (pRDA), conditioning each dimension of climate variation on the other two dimensions.

## RESULTS

### Alignment and Genotyping

Sequencing generated 404 million total reads across 189 individuals. We identified 150,710 SNPs through initial SNP calling, which were subsequently filtered to 10,760 high-quality SNPs, yielding an average sampling density of approximately one SNP per 42 kb. After filtering, we retained 175 individuals with a mean sequencing depth per individual of 40x (range: 11.7–111x). All retained individuals exceeded a minimum mean sequencing depth of 10x, ensuring that the dataset was of sufficient quality for downstream analyses.

### Population Structure and Genetic Diversity

Population structure and clustering patterns coincide with geography, riparian networks, and mountain ranges (Figure 1). Cross-validation error values from the ADMIXTURE analysis indicated optimal clustering of six distinct groups. For the purpose of visualizing population structure geographically, we plotted individuals by their majority ancestry from the ADMIXTURE analysis with K = 6 clusters (Figure 1C).

**Figure 1:**
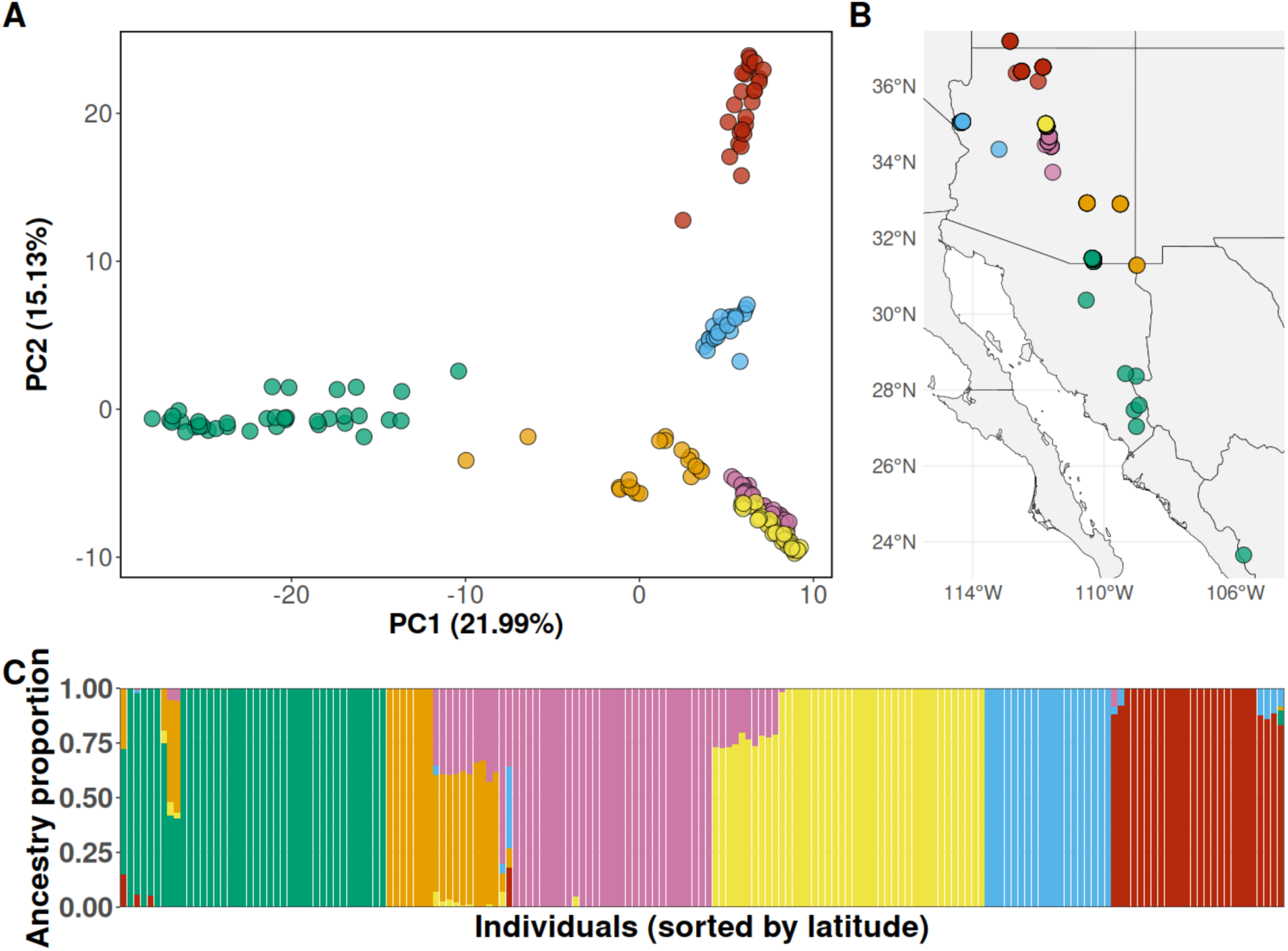
Crimson monkeyflower exhibits strong population structure and associations with latitude. **A** Individuals plotted by PC1 and PC2 from the PCA with each individual colored by majority ancestry from the ADMIXTURE analysis. **B** Individuals colored by majority ancestry from the admixture analysis and plotted on a map. **C** Results of ADMIXTURE analysis (K=6) where bars represent individuals and their proportion of ancestry organized by latitude increasing from left to right.

Fine scale population structure was observed within landscape features (mountain ranges and riparian corridors) with individuals forming distinct clusters based on sampling location within well sampled landscape features (Figure 2). Strong, identifiable population structure was observed in PCA space with tight clustering. Additionally, we see differentiation at extremely fine scales, including populations within 5km of each other.

**Figure 2:**
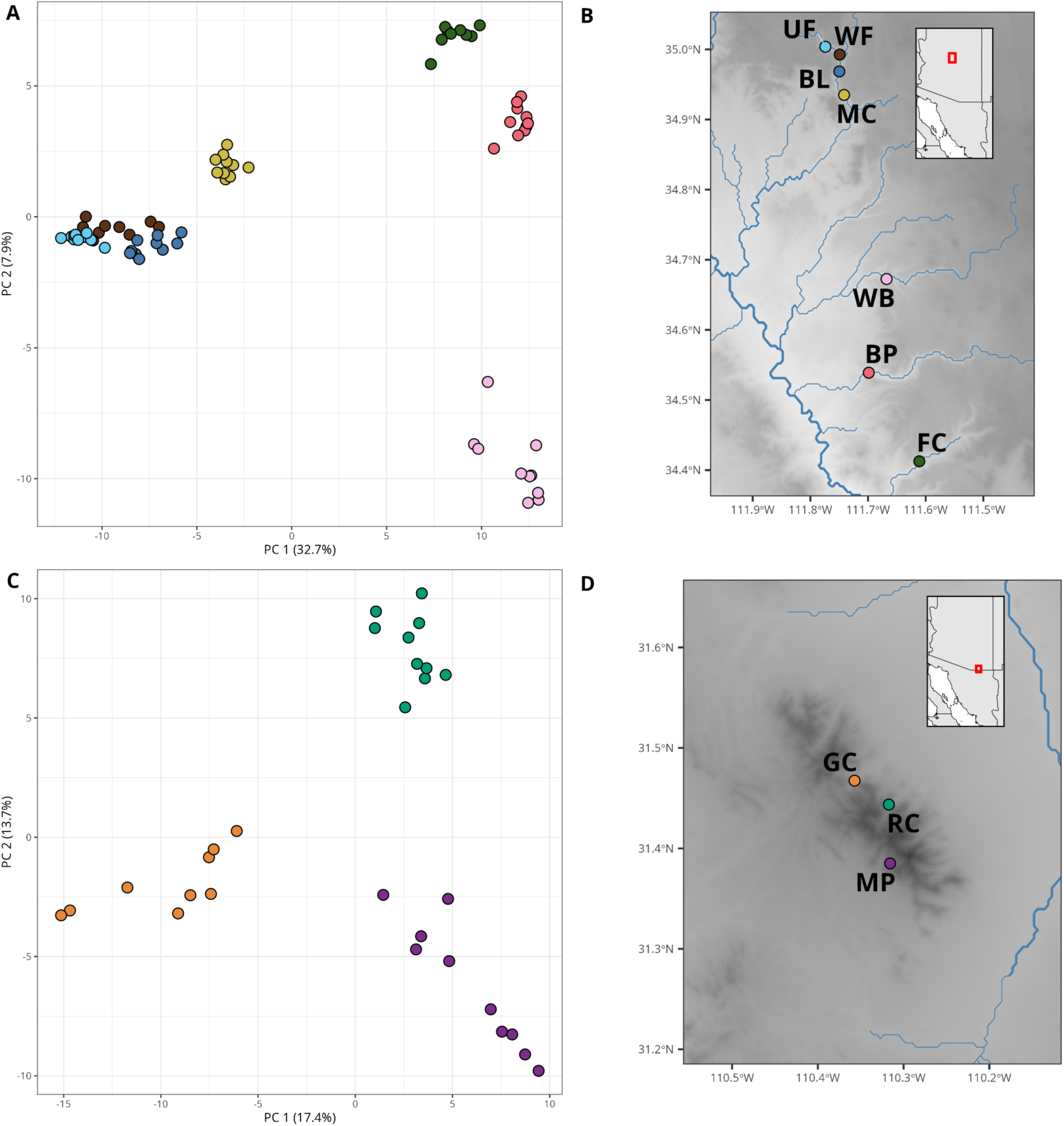
Fine scale population structure is observed in heavily sampled regions including a connected riparian corridor **(A and B)**, Verde Valley and Oak Creek (pink and yellow clusters from ADMIXTURE analysis [Fig. 1]), and a mountain range, Huachacas (green cluster from ADMIXTURE analysis [Fig. 1]), where populations are unconnected by perennial water **(C and D)**.

Consistent with findings of clustering based on landscape features, we found that genetic distance (*F*_ST_, Nei’s *D*) was lower between populations in the same riparian corridor or mountain range than it was between populations from different landscape features (Supplementary Figure 5). Population differentiation was two to ten times greater between populations from the same landscape feature than between populations within the same landscape feature. Pairwise *F*_ST_ between populations from the same landscape feature ranged from 0.032 to 0.146, while pairwise *F*_ST_ between populations from different landscape features ranged from 0.155 – 0.337. Pairwise Nei’s *D* between populations in the same region ranged from 0.032 to 0.147, while pairwise Nei’s *D* between populations in different regions ranged from 0.184 to 0.337. Levels of genetic diversity within populations were highly variable, with expected heterozygosity (H _e_) ranging from 0.052 to 0.18. Populations connected via perennial riparian corridors showed lower genetic diversity than populations in mountain ranges, all of which lacked riparian connectivity (Figure 3). Within riparian corridors, upstream populations consistently exhibited lower diversity than downstream populations.

**Figure 3:**
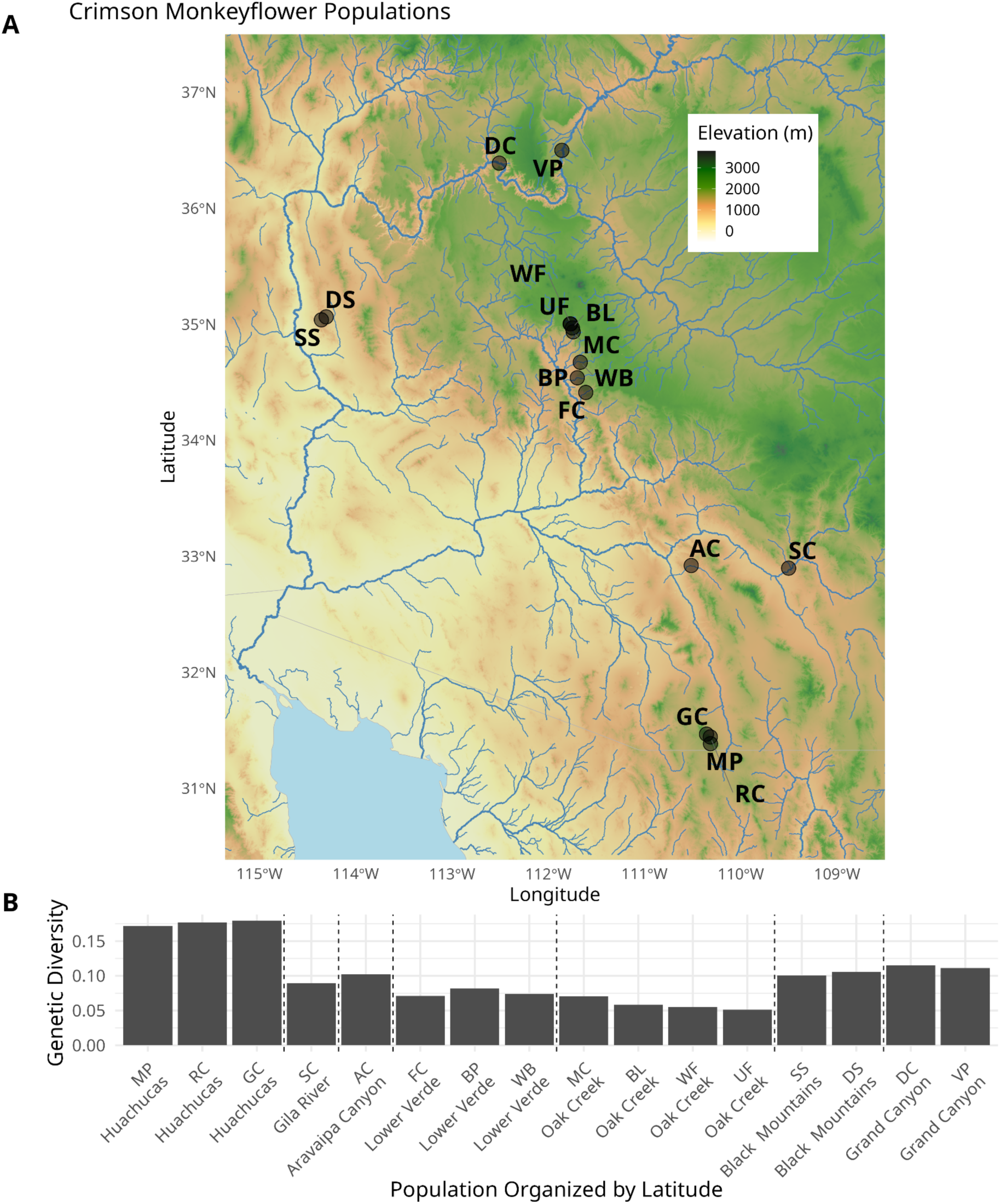
A. Sampling locations for population level data. All populations except DS, SS, GC, MP and RC have direct connectivity to riparian corridors via perennial water. **B.** Genetic diversity (expected heterozygosity) for each sampled population.

### Climate Variables Influencing Genetic Variation

After using PCA to reduce the dimensionality of climate variation, the first three dimensions of climate variation explained 96.4% of climate variation among our sites (Figure 4). A climate gradient ranging from warm, dry sites to cooler, wetter sites was the largest dimension of climate variation (Dim1). This dimension explained 61.9% of variation and was largely correlated to elevation (r = -0.80). As such, precipitation increases with elevation and temperature decreases with elevation across our sites. The second largest dimension of climate variability explained 17.5% of overall variation (Dim2). This dimension describes a gradient ranging from high summer precipitation and stable annual temperature to lower summer precipitation and higher annual temperature differences; this gradient is tightly correlated to latitude across sites (−0.96). Across sample sites, annual precipitation increased and continentality decreased as we moved from north to south. A gradient of relative humidity represented the third largest dimension of climate variation, explaining 12.6% of overall climate variation (Figure 4).

**Figure 4:**
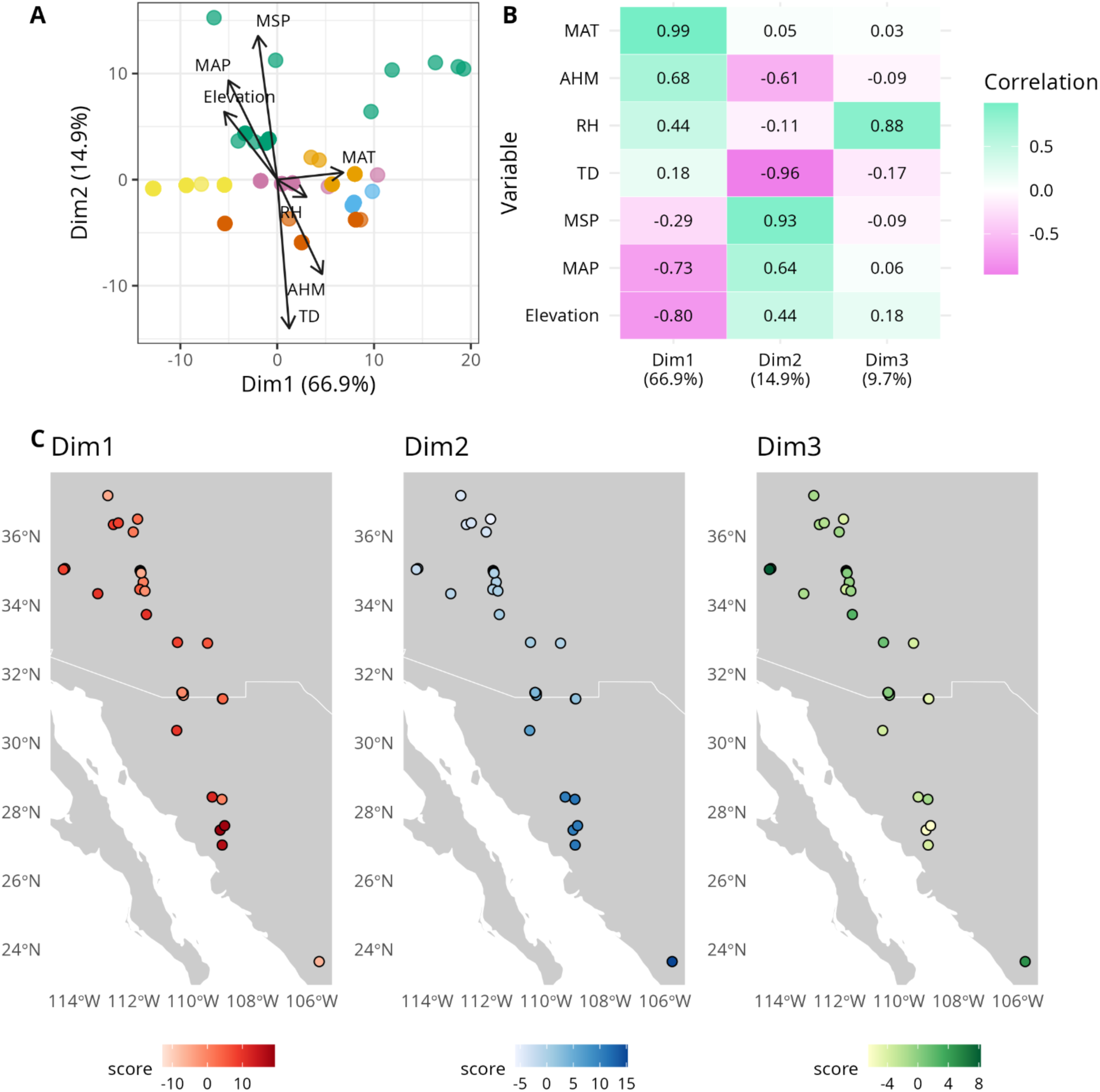
A. Shows a biplot of major climate variables with climate dimension 1 on the x axis and climate dimension 2 on the y-axis. Dots on the figure represent individuals in the climate space colored by their majority ancestry from the ADMIXTURE analysis. The correlation between each principal component and climate and geographic variables is shown in **B.** The relationship between each PC and geography is shown in **C**.

Geography and climate variation were both correlated with genetic variation in this species (Supplementary Figure 3). Mantel tests demonstrated that geographic distance and climate distance were both predictors of linearized *F*_ST_ (IBD: Mantel *r* = 0.576, *p* < 0.05; IBE: Mantel *r* = 0.499, *p* < 0.05). Geographic distance and climate distance were correlated to each other across the species range (Mantel r = 0.250, p < 0.05). Climate distance had a unique component that predicted genetic distance after controlling for geography (Mantel r = 0.471, p < 0.05). Some of the autocorrelation between geography and climate may be due to climate gradient Dim2 tracking closely with latitude across our species range. Yet other important dimensions of climate variation lack a correlation with latitude and instead track elevation (Dim1) or represent a relative humidity gradient independent of elevation, latitude, and most other climate variables (Dim3) (Supplementary Figure 5). These dimensions of climate variation (Dim1 and Dim3) vary strongly between nearby sites, allowing for local adaptation that is decoupled from broad scale geographic distance (Figure 4).

The individual contribution of each dimension of climate variation to genetic differentiation was demonstrated by our redundancy analysis. Our RDA showed that multiple aspects of climate variation contributed to genetic variation across our species range. The full RDA model explained 8.3% of total genetic variation with 53.3% of this variation explained by the first constrained axis and 32.0% of this variation explained by the second constrained axis (Figure 5). The pRDA showed that a gradient of temperature and annual precipitation (Dim1) predicted the largest unique fraction of variation at 2 .8%, while relative humidity (Dim3) predicted the second largest fraction at 2.1%, and summer precipitation with continentality (Dim2) predicted 1.7% of genetic variation (Figure 5).

**Figure 5:**
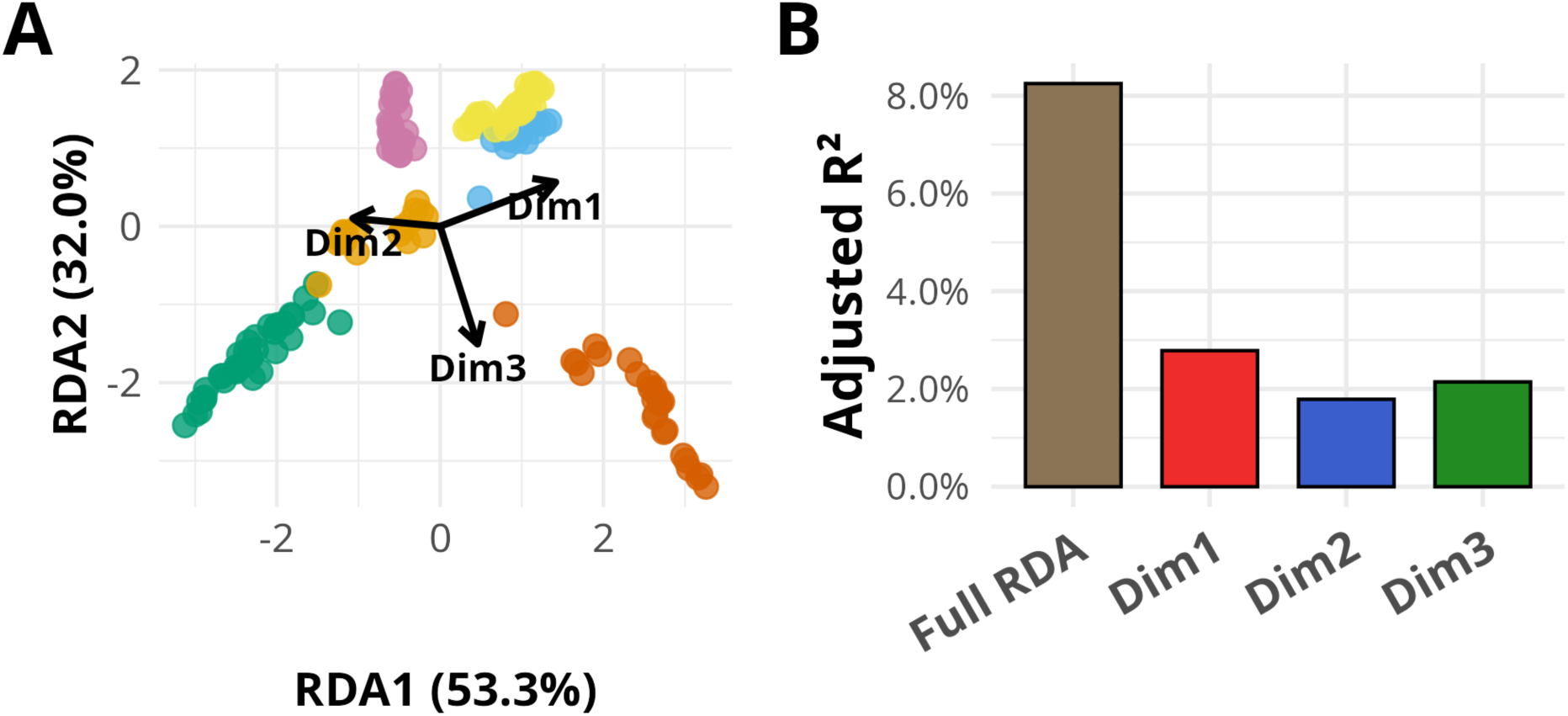
A. Axis 1 and 2 of the Redundancy analysis (RDA) of individuals’ genetic variation constrained by climate variation. Direction and length correspond to loadings of each dimension of climate variation on a given RDA axis. Colored dots correspond to majority ancestry from admixture analysis. **B.** Portion of genetic differentiation (Adjusted R ^2^) predicted by the full set of dimensions of climate variation (full RDA) and each dimension of climate conditioned on the other two dimensions (pRDA). The first dimension of climate variation (Dim1) explained the largest unique proportion of genetic differentiation. Despite representing the smallest dimension of climate variation Dim3 predicted the second largest proportion of genetic differentiation. Dim2 explained the smallest proportion of genetic differentiation of the three dimensions of climate variation.

## DISCUSSION

Our examination of a spring-specialist plant revealed patterns of diversity and differentiation distinct from those observed in most upland and generalist riparian plant species. Observations of population structure at remarkably fine scales likely represent the discrete nature of suitable habitat created by isolated springs. Additionally, patterns of diversity suggest that riparian connectivity may play a role in influencing genetic diversity locally within riparian corridors but not at broader scales. At broad scales, our results suggest that both demographic processes and local adaptation influence genetic differentiation in this species. While climate gradients driven primarily by precipitation are often important drivers of differentiation in upland plants, local adaptation in our species occurred along climate gradients that were more strongly associated with temperature, humidity, or continentality than precipitation alone.

### Population Structure and Genetic Diversity

Crimson monkeyflower populations show strong population structure across multiple spatial scales, influenced by both large scale landscape features such as mountain ranges and riparian networks, as well as fine scale structuring based on the individual springs that provide habitat for these plants (Figure 2). Fine scale genetic structure was detected across extremely small distances ( < 5km), both among populations connected by perennial riparian networks and among those lacking riparian connectivity within mountain ranges. Most sampling sites were represented by discrete pockets of habitat created by seeps and springs on canyon walls or isolated springs in an otherwise arid landscape. Strong population structure across small spatial scales has been observed in other spring-specialists including a spring-dependent water mite (*Partnunia steinmanni*), which exhibits spring specific genotypes (Blattner *et al*. 2022). The largely discontinuous distribution of spring-specialist species likely reduces gene flow and increases genetic drift within local populations, contributing to the strong local population structure observed.

One possible explanation for the presence of population structure at both broad and local scales is changes in connectivity of groundwater-dependent ecosystems due to post-Pleistocene drying across arid regions. This has been used to explain observations of population structure at multiple spatial scales in other spring-specialist species. Broad scale population structure likely represents ancient connectivity across the landscape while fine scale population structure is due to more recent isolation. Populations of relict dace ( *Relictus solitarius*), from nearby desert regions of North America, have broad population structure shaped by Pleistocene era pluvial lakes but fine scale structure correlated with isolated springs from more recent Holocene drying (Houston *et al*. 2015; Campbell *et al*. 2022). Similar patterns of both fine and broad scale population structure observed in the spring-specialist Kaiser newt (*Neurergus kaiseri*), endemic to the semi-arid Zargos Mountains of western Iran, may also be explained by recent climate oscillations, suggesting a broader pattern of recent differentiation among isolated spring-specialists (Goudarzi *et al*. 2019). Groundwater-dependent ecosystems across our species range experienced similar patterns of Pleistocene connectivity followed by Holocene drying as suggested by analyses of packrat middens in the region and pollen analysis from sediment cores (Baker 2008; Mitchell and Ober 2013; Galaz-Samaniego *et al*. 2023). Patterns of population structure observed in Crimson Monkeyflower align closely with those observed in spring-specialist animals and suggest a general pattern of fine scale population structure associated with recently isolated habitats.

At a local level, riparian connectivity influences genetic diversity. For populations within the same riparian corridor, upstream populations invariably exhibited lower genetic diversity than downstream populations (Figure 3). This finding is consistent with the unidirectional dispersal hypothesis, which predicts that upstream populations have lower genetic diversity than downstream populations when dispersal occurs via hydrochory (Anholt 1995; Honnay *et al*. 2010; Surmacz *et al*. 2025). Patterns of increased genetic diversity in downstream populations are strongest in pristine riparian corridors breakdown under habitat degradation, where negative impacts on downstream populations tend to be greater than those on upstream populations (Surmacz *et al*. 2025). Our ability to detect greater diversity in downstream populations in these riparian corridors demonstrates that these corridors retain sufficient connectivity to drive diversity gradients.

Across broad regional scales, riparian connectivity does not predict genetic diversity and frequently produces patterns opposite to the expectation of higher diversity within well-connected riparian corridors. Most sites nested within perennial riparian corridors showed lower genetic diversity than sites in mountain ranges lacking connectivity via perennial watercourses (Figure 3). Additionally, genetic differentiation between sites unconnected by riparian corridors was frequently lower than between sites connected by a riparian corridor. This suggests that at broad geographic scales, factors other than riparian connectivity play an important role in genetic differentiation. Once again, climate shifts since the end of the Pleistocene may explain this pattern; mountain ranges currently harbouring crimson monkeyflower populations may be acting as climate refugia for populations that were more continuous when lower elevations supported more suitable habitat.

### Isolation by Distance and Adaptation to Climate Gradients

A gradient from warm, dry sites to cooler, wetter sites was the largest climate influence on genetic variation in this plant (Dim1). Both temperature and precipitation are important drivers of local adaptation and genetic differentiation in plants (Whiting et al., 2024). Adaptation to this dimension of climate may be important in this species due to populations existing across a large elevational gradient (303m to 2685m) since temperature and precipitation are frequently correlated to elevation (Daly et al., 2008; Kelly & Goulden, 2008) . Elevation is also strongly correlated with this dimension, suggesting it captures the broad suite of climate variables underlying adaptation to this species’ large elevational range. Strong collinearity between variables makes it challenging to disentangle whether temperature or precipitation are driving genetic differentiation in this species. Previous research has shown that crimson monkeyflower exhibits tolerance to broader thermal conditions than sister lineages (Sheth and Angert 2014; Coughlin *et al*. 2022). This, combined with crimson monkey flower’s dependence on perennial water sources, which are often hydrologically decoupled from local precipitation, suggest that adaptation to different temperatures is an important driver of genetic differentiation in this species (Beardsley *et al*. 2003; Vickery 2008; Sheth and Angert 2014) .

Relative humidity was the second most important driver of genetic differentiation in crimson monkeyflower and was not found to be strongly collinear with other climate variables across our populations. This suggests humidity may represent a distinct selective gradient involved in structuring genetic variation across the species range. While climate gradients related to temperature and precipitation often dominate patterns of genetic variation, humidity can also drive adaptation in transpiration demand and stomatal conductance (Le *et al*. 2011; Tardieu *et al*. 2015; Driesen *et al*. 2020). Additionally, high humidity is the main predictor of fungal plant disease outbreaks and has the potential to increase adaptive pressures related to fungal pathogen resistance (Romero *et al*. 2022). High humidity not only increases the ability of fungi to colonize host plants (Li *et al*. 2014) but also reduces plant resistance to fungal pathogens by suppressing the activation of ethylene biosynthesis and signaling (Qiu *et al*. 2022). Atmospheric humidity may be an underappreciated climate variable for its influence on genetic variation, especially for spring-specialist plants living in environments where root moisture is at a near constant and is decoupled from short term changes in precipitation.

Summer precipitation and continentality also influenced genetic variation across this species. Summer precipitation across the crimson monkeyflower range is driven by monsoon patterns and is more intense and less predictable than other seasonal precipitation in the Sonoran Desert region (Scott *et al*. 2000). Since monsoon precipitation occurs during the hottest period of the year when plants’ water demands are highest, monsoon moisture may have greater importance for plant survival than moisture from other times of the year (Sponseller *et al*. 2012). Variation in monsoon precipitation has been suggested to drive seed dispersal timing in desert spineflower (*Chorizanthe rigida)* and growth patterns in creosote ( *Larrea tridentata)* (Sponseller *et al*. 2012; Martínez-Berdeja *et al*. 2015). This dimension also covaried with latitude, making it more challenging to disentangle with demographic processes such as isolation by distance than other dimensions of climate variation.

## Conclusion

Our results describe patterns of genetic variation in a species that spans a large elevational and latitudinal range but is restricted to habitats with perennial access to groundwater, primarily consisting of seeps and springs. Genetic variation in this spring-specialist plant is driven by an interplay of demographic and adaptive processes operating across multiple spatial and temporal scales. While riparian connectivity clearly influences local patterns of diversity and relatedness, its importance diminishes at broader regional scales where climate and geography act as stronger predictors of genetic differentiation. These patterns contrast with purely riparian and upland plant species, which exhibit more continuous variation at local scales. This pattern mirrors that found in spring-specialist animal taxa in North American deserts caused by ancient connectivity during wetter Pleistocene conditions and more recent isolation driven by post-glacial aridification. The broad landscape features it inhabits, canyon bottoms and mountain ranges, represent refugia from drier Holocene conditions, while seeps and springs within these sites provide further stability decoupled from short term and regional drought. As anthropogenic climate change threatens to make climate in this region hotter, drier, and less predictable at a rate never seen in recent ecological time, it remains to be seen if groundwater-dependent ecosystems will continue to serve as viable refugia. Understanding these ecosystems, their foundational plant species, and the patterns of diversity and differentiation in spring-specialists will be essential for informing and monitoring efforts to protect these rare and valuable ecosystems.

## SUPPLEMENTARY DATA

Supplementary data are available at *Annals of Botany* online and consist of the following. A supplementary dataset which contains: Table S1: Sampling locations for populations and herbariam samples. Table S2: Definitions of ClimateNA variables used in analysis and associated units. Table S3: Annual ClimateNA variables for each sample site. Table S4: Seasonal ClimateNA variables for each sample site. Table S5: Pairwise correlations among climate variables. Figure S1: Population structure inferred using ADMIXTURE for K = 2-10. Figure S2: Cross-validation errors for ADMIXTURE models K = 2-10 used to evaluate support for numbers of genetic clusters. Figure S3: Pairwise genetic differentiation among populations. Figure S4: Relationship among genetic, geographic and climate distance.

## FUNDING

This research was partially supported by the Northern Arizona University (NAU) Research and Creative Activity Support Grant to L.M.H.

## CONFLICTS OF INTEREST

The authors declare no conflicts of interest.

## AUTHOR CONTRIBUTIONS

M.W. and L.M.H. conceived and designed the research. L.M.H. supervised the research and secured funding. M.W. completed tissue collection and lab work. M.W. performed bioinformatics and data analysis with assistance from T.M.F. M.W. wrote the original manuscript. All authors reviewed and edited the draft manuscript and approved the final manuscript.

## AVAILABILITY OF DATA AND MATERIALS

The data that support the finding of this study will be made available on the Dryad Digital Repository upon publication. Scripts for analysis will be made available on Github or Dryad Digital Repository upon publication.

## ACKNOWLEDGMENTS

We thank Rob Massatti and Erika Sukovich for assistance with RADseq library preparation. Our thanks go to Emily Palmquist, Arturo Castro, Neal Javenkoski, Cassidy Phoenix, Brenda Weiss, and Thom Weiss for help with tissue collection. We are grateful to Betsy Black for helping obtain climate data. We thank Yaowu Yuan for the *M. verbenaceus* genome. Tissue collection in Grand Canyon National Park was conducted under permit No. GRCA-2019-SCI-0052.

## Notes

### Competing Interest Statement

The authors have declared no competing interest.

