## Supplementary Figures for "Climate Gradients and Habitat Discontinuity Structure Genetic Variation in a Spring-Specialist Plant"

Matthew Weiss[1], Trevor M. Faske[2], Liza M. Holeski[1]
[1]Department of Biological Sciences, Center for Adaptable Western Landscapes, Northern Arizona University, Flagstaff, Arizona, USA [2]Landscape Stewardship Collective, Flagstaff, AZ, USA

### Table of Contents:

**Figure S1.** Population structure inferred using AMIXTURE for K = 2-10. Bar plots show individual ancestry proportions across inferred clusters. The main text presents K = 6.

**Figure S2.** Cross-validation errors for ADMIXTURE models K = 2-10 used to evaluate support for numbers of genetic clusters.

**Figure S3.** Pairwise genetic differentiation among populations. A heatmap showing pairwise $F_{ST}$ (lower triangle) and Nei's *D* (upper triangle) among sampled populations

**Figure S4.** Relationship among genetic, geographic and climate distance. Scatter plots showing isolation by distance (IBD), isolation by environment (IBE), spatial autocorrelation of climate (geographic vs. climate distance), and IBE conditioned on geographic distance.

**Figure S1. Population structure inferred using ADMIXTURE for K = 2-10.**

Bar plots show individual ancestry proportions for each inferred number of genetic clusters (K). Each vertical bar represents an individual, and colors correspond to inferred ancestry components. Individuals are organized by latitude from south to north. The main text presents results for K = 6.

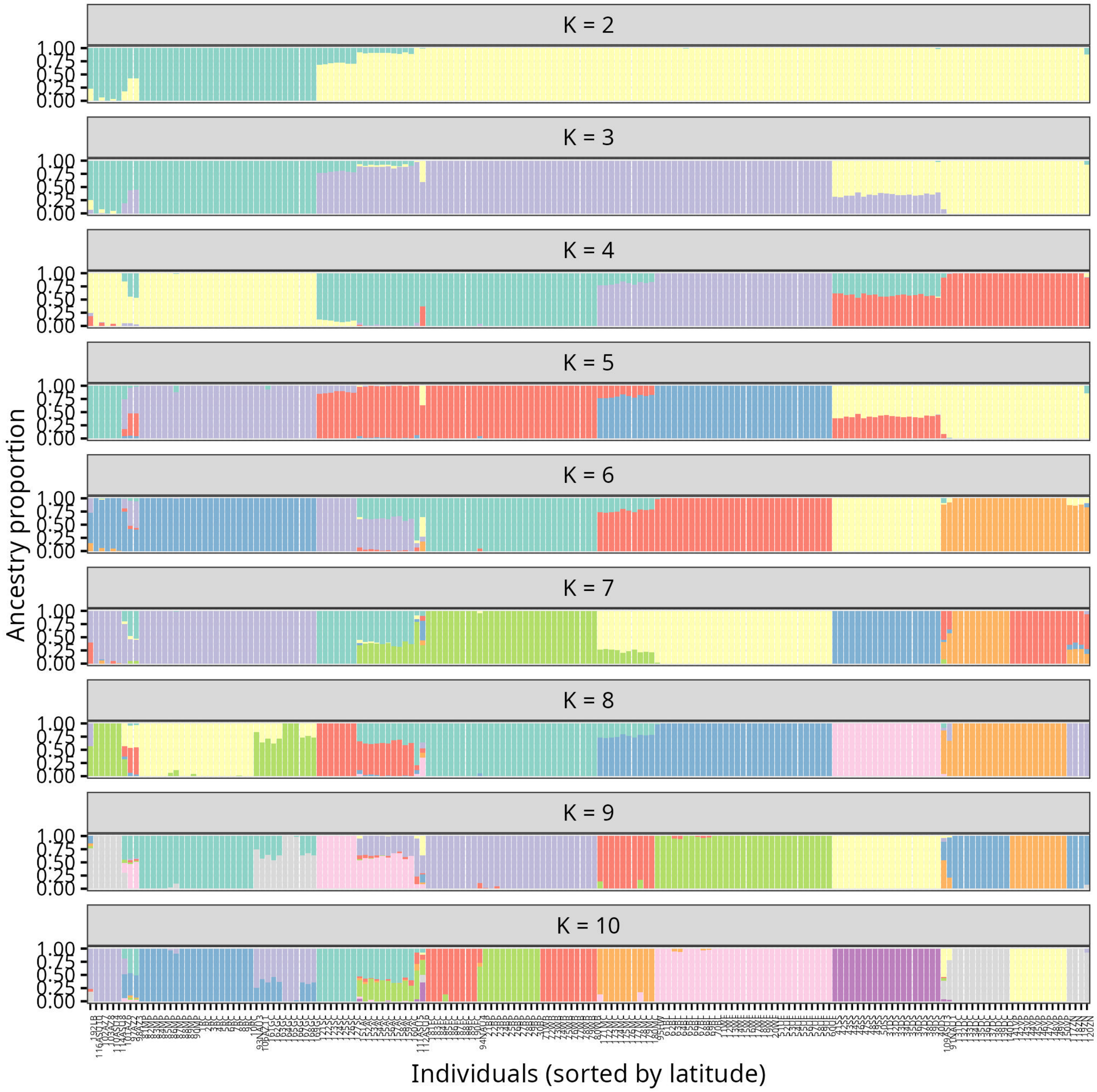

**Figure S2. Cross-validation error across ADMIXTURE models.**

**A** Cross-validation (CV) error values for K = 2-10. Lower CV error indicates improved model support. **B** Change in CV error between K values (−ΔCV error). Larger values in −ΔCV error indicated greater improvement in model fit with increasing K.

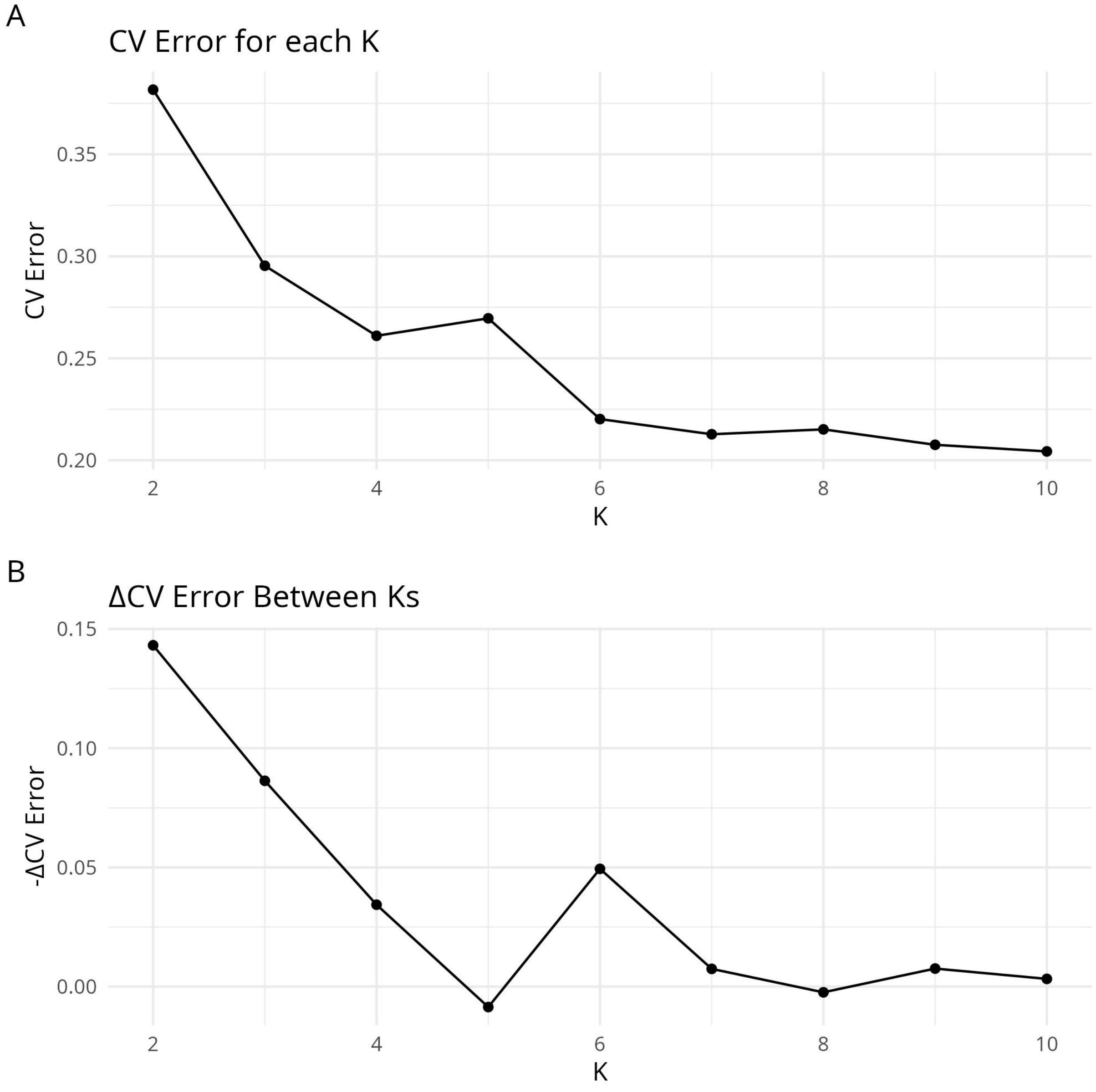

**Figure S3. Pairwise genetic differentiation among populations.**

Heatmap showing pairwise genetic differentiation among populations. Populations are grouped by latitude and black lines separate populations from different riparian networks or mountain ranges. The lower triangle displays $F_{ST}$ values, and the upper triangle shows Nei's genetic distance (Nei's *D*). Color intensity reflects the magnitude of genetic differentiation.

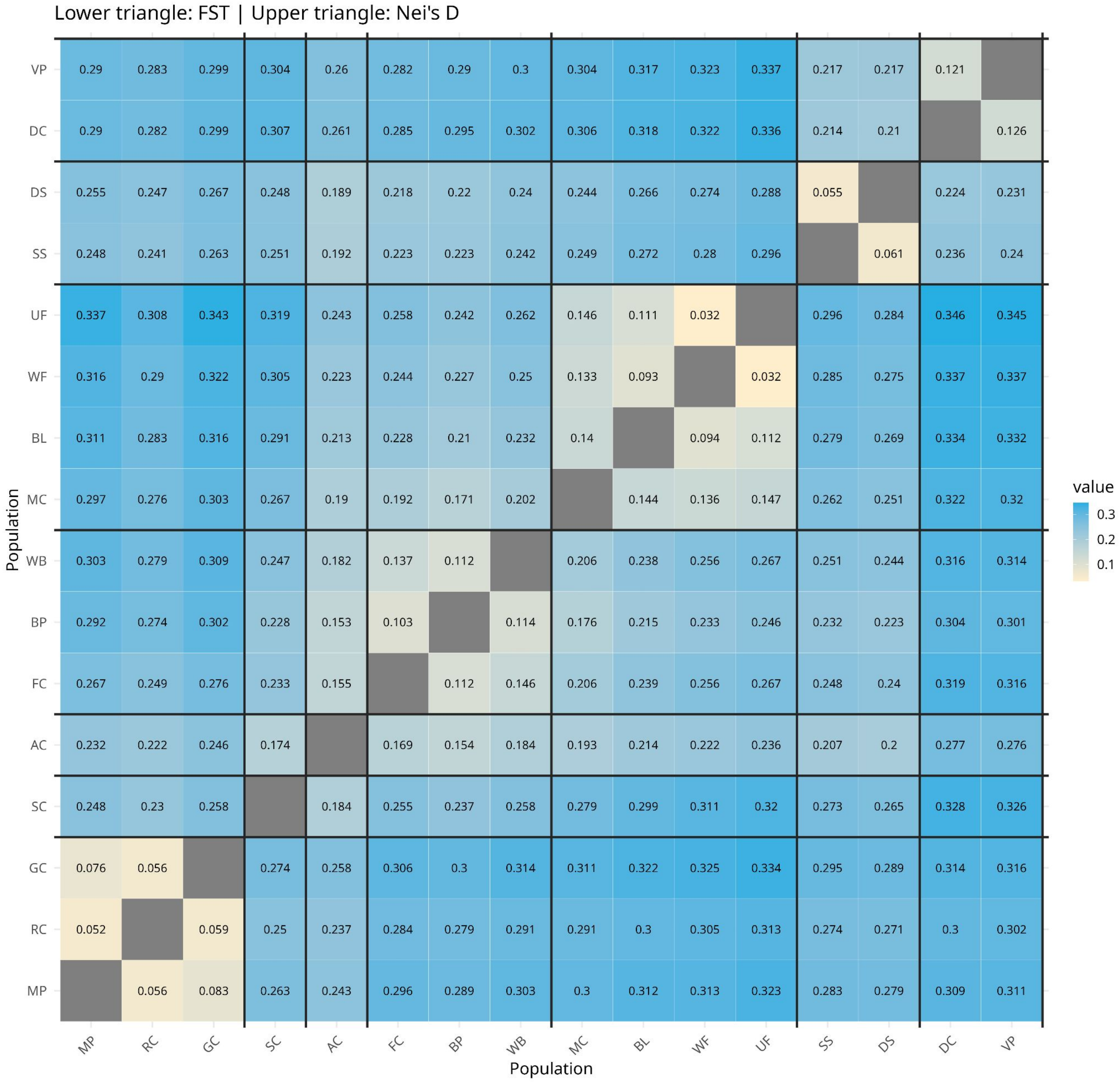

**Figure S4.** **A** Mantel test showing the relationship between pairwise geographic distance and pairwise genetic distance (linearized $F_{ST}$; $F_{ST}$ / (1-$F_{ST}$) ). **B** Mantel test showing the relationship between pairwise climate distance and genetic distance (linearized $F_{ST}$; $F_{ST}$ / (1-$F_{ST}$) ). **C** Mantel test showing the relationship between pairwise climate distance and genetic distance (spatial autocorrelation of climate variation). **D** Partial Mantel test showing the relationship between pairwise climate distance and genetic distance (linearized $F_{ST}$; $F_{ST}$ / (1-$F_{ST}$) after controlling for geographic distance.

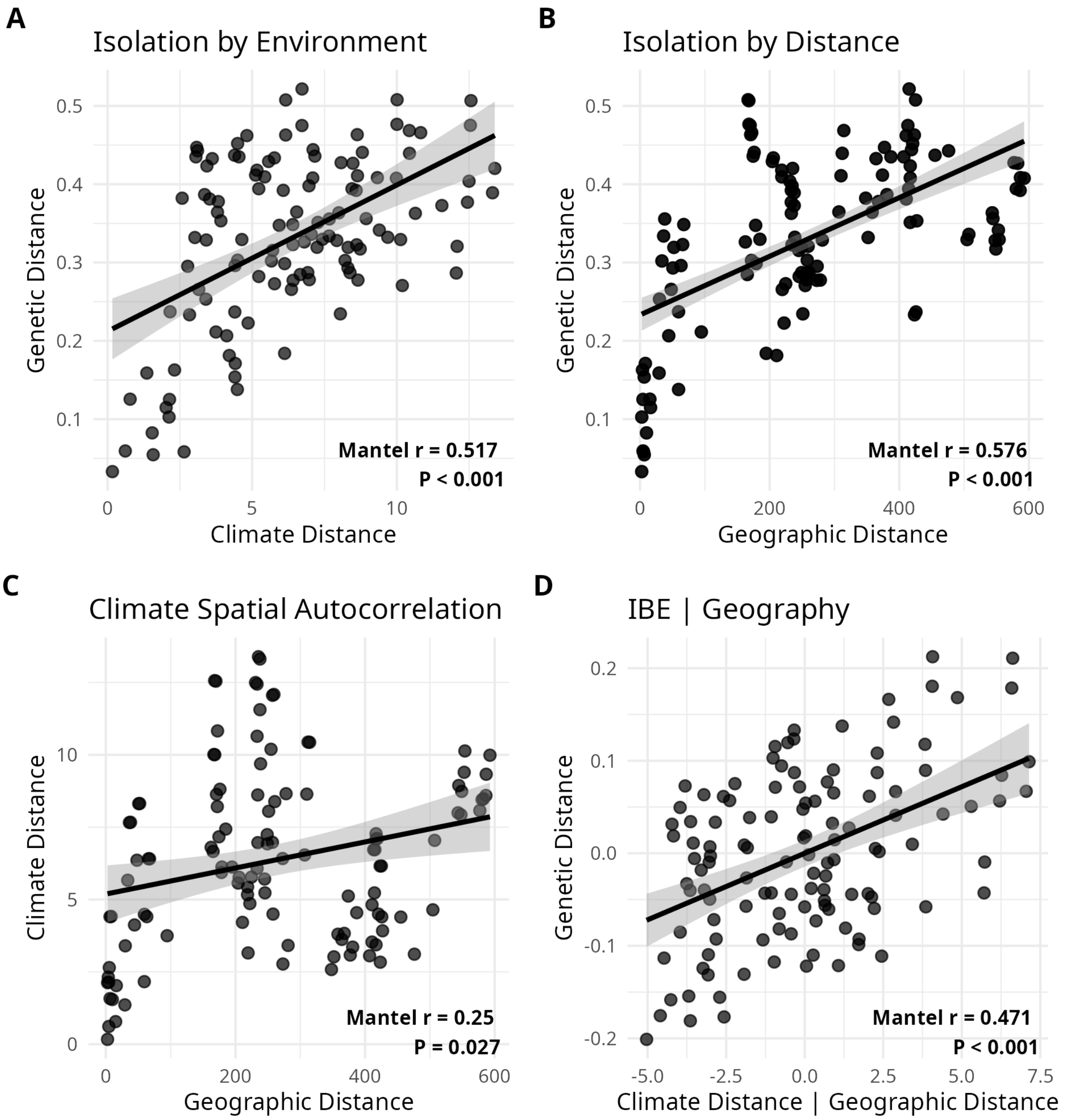
